# Evasion of Toll-like Receptor Recognition by *Escherichia coli* is mediated via Population Level Regulation of Flagellin Production

**DOI:** 10.1101/2021.11.18.469103

**Authors:** M Lanz, C Birchall, L Drage, D Picton, C Mowbray, Q Alsenani, A. Tan, A Ali, C Harding, R Pickard, J Hall, PD Aldridge

## Abstract

Uropathogenic *Escherichia coli* (UPEC) is a major cause of urinary tract infections. Analysis of the innate immune response in immortalised urothelial cells suggests that the bacterial flagellar subunit, flagellin, is key in inducing host defences. A panel of 40 clinical uro-associated *Escherichia coli* isolates recovered from either asymptomatic bacteruria (ASB), cystitis or pyelonephritis patients, were characterised for motility and their ability to induce an innate response in urothelial cells stably transfected with a NFκB luciferase reporter. Twenty-four isolates (60%) were identified as motile with strains recovered from cystitis patients exhibiting a bipolar motility distribution pattern (P < 0.005) and associated with a 2-5 fold increase in NFκB signalling. Although two isolates were associated with swarm sizes of >7 cm and NFκB activities of >30 fold (P = 0.029), data overall suggested bacterial motility and the NFκB signalling response were not directly correlated. To explore whether the signalling response reflected antigenic variation flagellin was purified from 11 different isolates and the urothelial cell challenges repeated. Purified flagellin filaments generated comparable (30.4±1.8 to 46.1±2.5 fold, P = NS) NFκB signalling responses, irrespective of either the source of the isolate or H-serotype. These data argued against any variability between isolates being related to flagellin itself. To determine the roles, if any, of flagellar abundance in inducing these responses flagellar hook numbers of a range of cystitis and ABU isolates were quantified using a plasmid encoded flagellar hook gene *flgEA240C*. Foci data suggested isolates were averaging between 1 and 2 flagella per cell, while only 10 to 60% each isolates population exhibited foci. These data suggested selective pressures exist in the urinary tract that allow uro-associated *E. coli* strains to maintain motility exploiting population heterogeneity to prevent host TLR5 recognition.

## Introduction

Urinary tract infections (UTIs) are among the most common bacterial infections suffered by individuals of all ages. They affect an estimated 150 million people worldwide including children, young adults and older populations (1). Infections are often painful and debilitating, associated with a wide range of pathogens, but the majority (70-80%) link to the bacterial uropathogen *Escherichia coli* (2). Regardless of the uropathogen, treatment options remain limited with antibiotics being the first choice therapeutic. Treatment consequences, namely multi-drug resistant bacteria, often underpin persistent or rUTIs and has driven the urologic community to work collaboratively to adopt antibiotic stewardship programmes (3)

Research to date suggests UTIs link to genotypic and phenotypic variation in both the host and the uropathogen (4,5). At present it is assumed that the relationship between an individual’s susceptibility and bacterial virulence determines the balance between tolerance of invading pathogens and the mounting of an immune response, which in turn dictates the course of infection and subsequent recurrence (6–8). *Escherichia coli* reside naturally in the gastrointestinal tract, but are able to migrate from the anus, colonise the vaginal and periurethral areas, then ascend to the bladder causing asymptomatic infection (ABU) or acute cystitis (5). However, our understanding of the associated host-microbe interactions is compounded by the observation that the same or related strains can lead to both symptomatic UTI and ABU. One outcome is that ABU patients, particularly the elderly, are often given antibiotics without justification due to clinical uncertainty (1).

While UPEC harbour a large array of virulence determinants, the ability to cause disease is dependent on the ability of the bacterium to ascend the urinary tract through adherence (fimbriae driven) and flagella-based motility (2). Moreover *in vivo* studies using genetically engineered UPEC strains and mice UTI models support flagella as being a key factor in the aetiology of an UTI (9–11). The bacterial flagellum is a macromolecular, self-assembling nano-machine whose genetics, assembly process and mechanisms of action during host-microbe interactions are well-documented (12–17). *E. coli* is known to produce 2-8 flagella per cell arranged peritrichously across the cell surface and, is characterised genetically, by approximately 60 flagellar genes organised into three loci: *flg*, *flh* and *fli* that function to orchestrate flagellar assembly and rotation (18). Evidence supports flagellar assembly and function to be coupled to flagellar gene expression by a complex transcriptional hierarchy (19). Additionally, tight control of flagellar gene expression enables *E. coli* to efficiently pass-through ON/OFF phases of motility that can be exploited and used advantageously during host-microbe interactions (20).

In humans, uropathogens such as UPEC are sensed via TLR5 receptors, which detect flagellin: the major subunit of the flagellum filament (21). TLR5 activation results in the rapid release of urothelial host defence agents including cytokines and defensins that function individually or collectively to kill potential uropathogens (22,23). However, using urine and employing *in vitro* chemotaxis assays Herrmann and Burman (1985) reported that only 68% (19/28) of *E. coli* isolates associated with cystitis, or an UTI, were motile (24). Yet, there is strong evidence to support uropathogenic *E. coli* (UPEC) exploiting flagellar-mediated movement to establish the initial ascending colonisation of the bladder from the urethra (11,25). Lane et al (2005) and Wright et al (2005) both concluded that motility provided UPEC a competitive advantage over non-motile UPEC strains in establishing an UTI in murine models. A key challenge therefore is to understand what triggers potential uropathogenic bacteria to regulate their motility, ascend the urethra and infect the bladder.

Using clinically derived uro-associated *E. coli* isolates data are presented suggesting a regulatory mechanism linked to population heterogeneity that maintains motility within a bacterial population, but at levels below a threshold required for innate immune recognition.

## Results

### Uro-associated E. coli motility and the urothelial innate response

Forty uro-associated *E. coli* isolates were curated from patients presenting with either cystitis, pyelonephritis, asymptomatic bacteriuria (ABU) or UTI-associated bacteraemia. Semi-quantitative agar assays measuring the size of a bacterial swarm after 8 hrs (Figure 1A) were used to assess motility of these isolates and 24 (60%) were identified as motile (Fig 1A & B, Table S1). In general, the swarms of isolates recovered from ABU patients measured between 0.8 and 5.4 cm, while cystitis strains exhibited a bipolar motility distribution pattern with strains swarming less (n=6) or greater (n=4) than 5.4 cm respectively (Fig 1B; P < 0.005).

**Figure 1:**
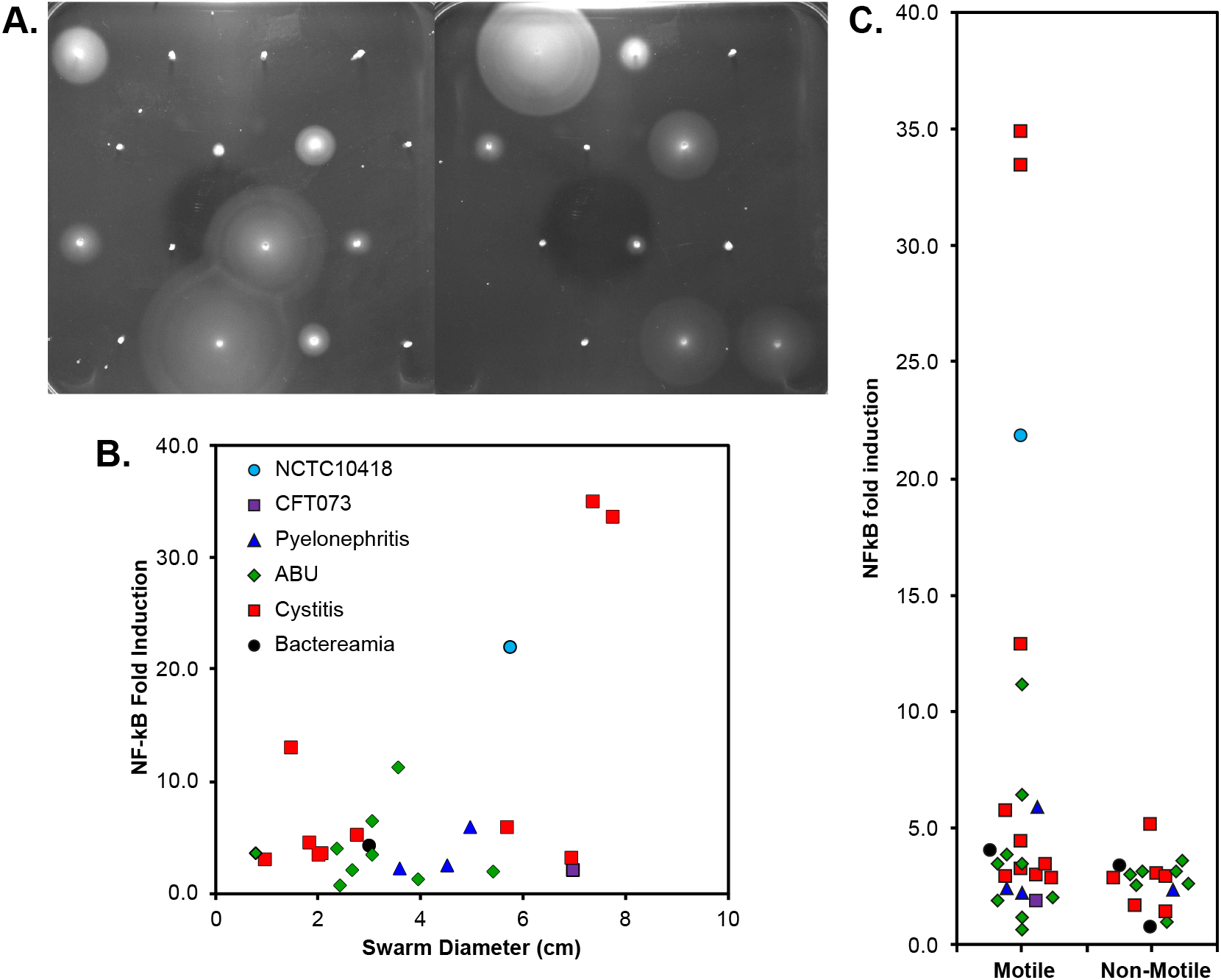
Motility of clinical uro-associated isolates and RT4 bladder cell NF-κB signalling. **A)** Motility agar assay data of a selection of clinical UPEC isolates (Table S1 and S2) showing the diverse range of phenotypes. **B)** Quantification of swarm diameter for n=3 independent colonies of each clinical isolate plotted against NFκB induction. Control strains, NCTC10418 and CFT073, and clinical isolates are colour coded. **C)** NFκB fold induction of all motile and non-motile UPEC clinical isolates. Points have been scattered left or right with respect to the x-axis for clarity.

The impact of bacterial motility on the urothelial innate response was assessed *in vitro* using heat-killed isolates (1 × 10^5^ CFU/ml) and bladder RT4 cells stably transfected with a NFκB luciferase reporter (26). Following these challenges 33 (83%) of all the isolates were associated with a 2-5 fold increase in NFκB signalling (Fig 1B & C), with 7 of the motile isolates associated with increases of >5-fold. Two isolates recovered from cystitis patients were associated with NFκB activities of >30 fold (Fig 1C; P = 0.029) and swarm sizes of >7 cm (Fig 1B). These data suggested that bacterial motility linked to a NFκB signalling response, but that the two were not directly correlated.

### Urothelial responses to flagellins prepared from uro-associated E. coli isolates

UTIs are ascending infections and urothelial cells respond to potential uropathogens via flagellin detection, TLR5 signalling and the release of antimicrobial killing and pro-inflammatory agents (26). TLR5 proteins recognise a conserved motif found in the majority of flagellins (27), which in *E. coli* is referred to as the H-antigen with to date 53 flagellin or H-serotypes being identified (28) (Table S1). To explore whether the signalling response reflected antigenic variation flagellin was purified from 11 different uro-associated *E. coli* isolates (Table S1: Lab IDs PYL3398-PYL3424) and the urothelial cell challenges repeated (Fig 2A). Purified flagellin filaments generated robust and comparable (30.4±1.8 to 46.1±2.5 fold, P = NS) NFκB signalling responses, irrespective of either the source of the isolate or H-serotype. These data argued against any variability between isolates being related to flagellin itself.

**Figure 2:**
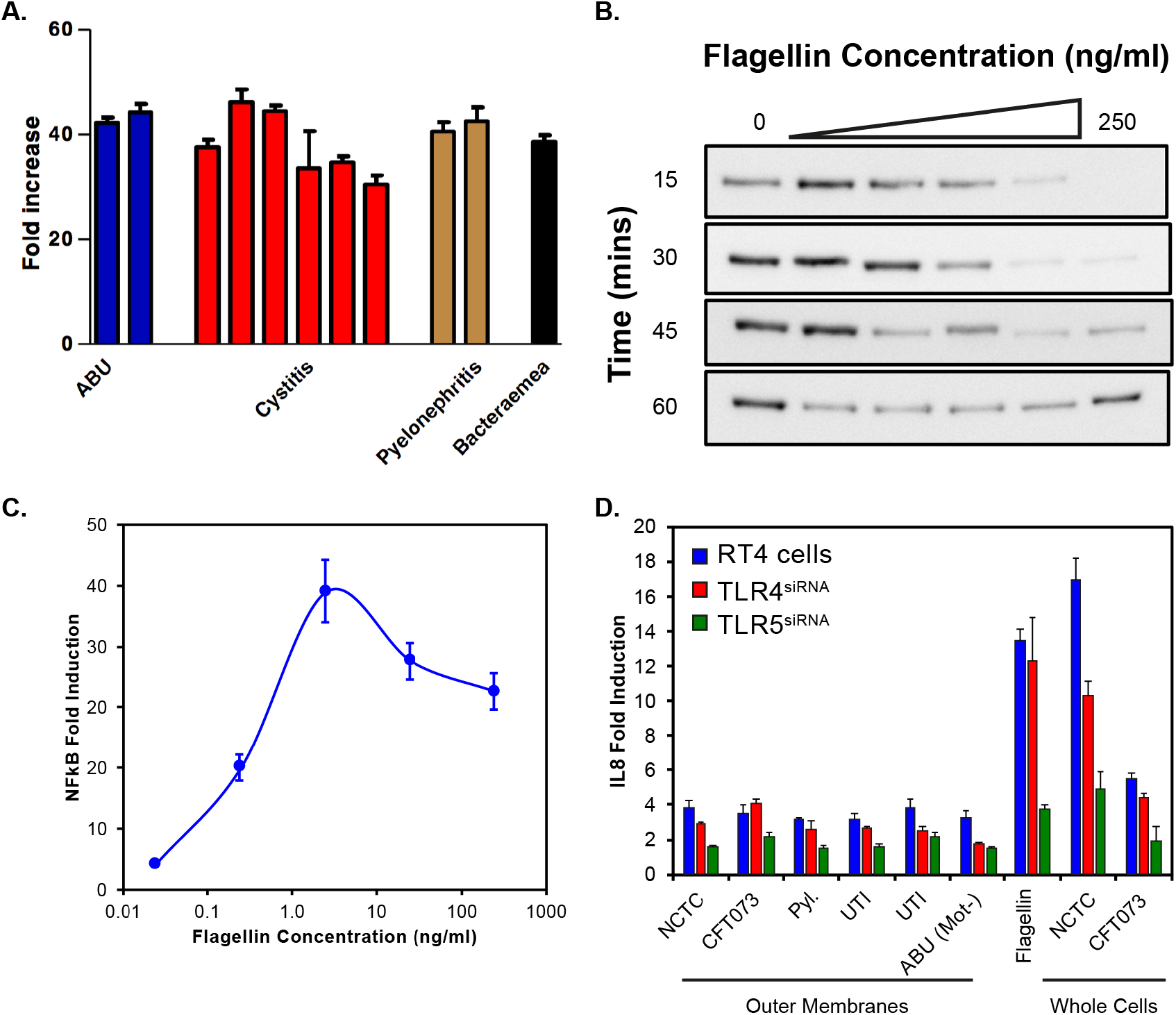
NF-κB and IL-8 responses of bladder RT4 cells challenged with flagellin, bacterial outer membrane preparations and whole cells. **A)** NF-κB response of RT4 bladder cells challenged for 24 hours with flagellin filaments (250 ng/ml) isolated from 11 different uro-associated *E. coli* (strains PYL3398-PYL3324: Table S1). **B)** Immunoblot analysis of IκBα following dose dependent challenges of RT4 cells with flagellin filaments isolated UTI3408. **C)** Quantification of NFκB induction following dose dependent challenges of RT4 cells with flagellin filaments isolated from UTI3408. Data shown is the average of two technical repeats and 3 independent challenges. **D)** IL-8 concentrations of RT4 cell media following transfection with siRNAs targeting either TLR4 or TLR5 expression and challenging for 24 hours with either outer membrane preparations, flagella filaments (50 ng/ml) or heat killed whole bacteria.

TLR-signalling activates a complex signalling cascade that leads to the degradation of IκBα and NFκB release that activates gene expression (29). Smith et al (2003) using transfected Chinese hamster ovary cells, showed TLR5 recognition of flagellin to be dose-dependent (30). A dose-dependent response to flagellin, measured through IκBα protein levels and NFκB induction was also observed in RT4 urothelial cells (Fig 2B and C). Interestingly, using purified flagelllin from *E. coli* UTI3408 experiments suggested a peak response at approximately 2.5 ng/ml.

### Urothelial responses to outer membranes prepared from uro-associated E. coli isolates

Data from *in vitro* and *in vivo* studies, both murine and clinical, suggest TLR4 signalling as well as TLR5 impacts UTI host-microbe interactions (21,31–33). TLR4 recognises lipopolysaccharide, the major outer membrane (OM) component of Gram-negative bacterial species such as UPEC (34). Therefore, the potential roles of TLR4 and LPS were explored to help explain the differing NFκB signalling responses observed during whole bacterial cell challenges. Outer membrane preparations of four clinical isolates (3 motile [PYL3398, UTI3408 and UTI3412] (Table S1) and 1 non-motile [ABU3416]) and two control strains NCTC10418 and CFT073, were used to challenge RT4 (TLR^+^) cells and RT4 cells where either TLR4 or TLR5 expression had been inhibited by siRNA knockdown (Fig 2D). The innate response was determined through measurement of the pro-inflammatory cytokine IL-8. Data presented in Fig 2D showed that the IL-8 responses to flagellin (50 ng/ml) and whole bacterial cell challenges were significantly reduced in the TLR5^siRNA^ cells (P = 0.0013). While a two-fold reduction in IL8 was observed in the NCTC14028 challenged TLR4^siRNA^ cells. This decrease was not significant and not mirrored in either the flagellin or CFT073 challenges. Challenges with OM sample preparations supported reduced IL-8 concentrations overall, but again a significant reduction was detected only in the TLR5 silenced cells (Fig 2D P < 0.001). While these data suggested the OM preparations maybe contaminated with flagellin they also supported minor roles for LPS and TLR4 in the RT4 bladder cell innate response to an acute infection.

### Correlating urothelial responses to flagellar abundance amongst uro-associated E. coli isolates

Data in Fig 1 and 2 indicated that flagellin isolated from a range of uro-associated *E. coli* induced an innate response as shown by NFκB signalling and effector (IL-8) responses. To determine the roles, if any, of flagellar abundance in inducing these responses flagellar hook numbers of cystitis (5), ABU (10) and PYL (1) isolates were quantified (Fig 3A) using the plasmid encoded flagellar hook gene *flgEA240C* (20). The control *E. coli* strain NCTC10418 averaged 2.5 foci per cell, while foci data suggested the clinical isolates and CFT073 were averaging between 1 and 2 flagella per cell (P = 0.148) (Fig 3B, x-axis).

**Figure 3:**
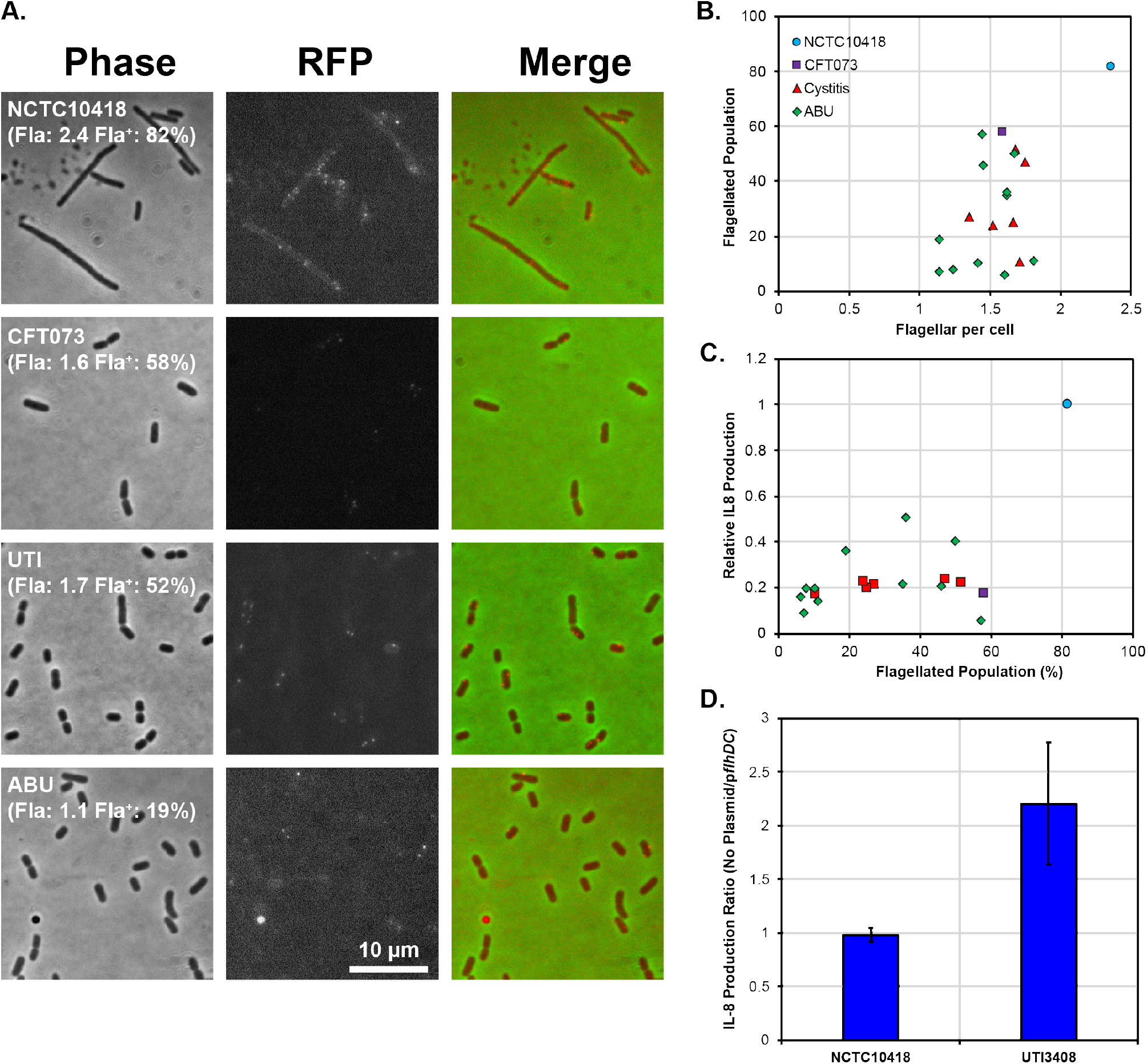
Uro-associated *E. coli* population heterogeneity and UPEC evasion of the TLR5 response. **A)** Phase contrast and fluorescent images of FlgEA240C foci in the control strains NCTC10418 and CFT073, and two uro-associated clinical isolates. Quantification of a minimum of 250 cells per strain is shown in brackets where Fla: = average number of FlgE foci per cell and Fla+: = percentage of the population with foci. Images are chosen to show foci and may not reflect quantified numbers. **B)** Scatter plot showing average foci per cell versus flagellated population. The range of average foci per cell for all strains except NCTC10418 are within experimental error (P = 0.143). **C)** Scatter plot showing percentage of flagellated population versus relative NFκB induction (control strain NCTC10418 = 1.0). **D)** IL-8 production, presented as a ratio, following challenges of RT4 cells with NCTC10418 (p*flhDC*-ve) or UTI3408 (p*flhDC* transformed).

However, only 10 to 60% of the CFT073 and clinical isolate cell populations exhibited FlgEA240C foci compared to 80% of the NCTC10418 cell population (Fig 3B, y-axis). Moreover, using NCTC10418 as the control and exploiting IL-8 concentrations resulting from RT4 urothelial cells challenge experiments, identified a link between flagellar numbers and the host response (Fig 3C P = 0.013). These data suggested selective pressures exist in the urinary tract that allow uro-associated *E. coli* strains to maintain motility but exploit population heterogeneity to prevent host TLR5 recognition and bacterial killing.

The flagellar system is regulated at the expression level by a transcriptional hierarchy controlled by the master transcriptional regulatory FlhD_4_C_2_ (35). FlhD_4_C_2_ levels are sensitive to a wide range of regulatory mechanisms that include transcription, translation and protein stability (36–39). Population data (Fig 3B) suggested that the number of flagella observed linked to reduced flagellar gene expression. To examine this further a high copy number plasmid encoding *flhDC* was transformed into the control strain NCTC10418 and UTI3408 (25% Fla+; 1.66 Fla/cell; 0.19 relative IL-8 production: Table S1). RT4 urothelial cell challenge experiments performed using these transformed strains showed that increasing *flhDC* expression supported a two-fold increase in IL-8 concentrations for UTI3408 compared to no change for NCTC10418 (Fig 3D).

## Discussion

Motility is a well-recognised pathogenicity, virulence and/or colonisation factor for a wide range of bacterial species including uropathogenic *E. coli* (UPEC) (40). However, motility links to flagellin production, which is the bacterial ligand for the mammalian host receptor TLR5. TLR5 activation releases host bacterial killing agents including cytokines and antimicrobial agents that facilitate bacterial killing, and clearance from potential colonisation and/or infection sites (21,26,30). Data from this study exploiting clinical isolates associated with UTIs suggest that uro-associated *E. coli* exploit population heterogeneity to maintain motility, but prevent the TLR5-dependent activation of the host innate immune response (41). Essentially these bacterial populations manipulate their flagellar production so they can survive and/or colonise the urinary tract, but remain under the host radar. It has been reported that *E. coli* swims efficiently with only one flagellum per cell (42) and using a FlgEA240C foci labelling approach uro-associated clinical isolates and the UPEC model strain CFT073 cultured *in vitro* were characterised by 1 to 2 flagella per cell (Fig 3). Together these data support *E. coli* motility and survivability within the lower urinary tract environment. However, once motility is enhanced through increased flagellar production, modelled *in vitro* using strain 3408 (Fig 3D), these microbes become visible to urothelial cell TLRs. Detection is associated with the release of a plethora of host defence molecules, shown here in these studies by elevated IL-8 concentrations (Fig 3D).

Flagellar systems have been shown to be subject to multiple regulatory controls (43). For example, *Salmonella enterica* generates a bipolar Fla^−^/Fla^+^ population in response to either nutritional and/or cell envelope stresses (44) while *Caulobacter crescentus* divides asymmetrically to produce one motile cell each division, ensuring a subpopulation of motile progeny within a growing population (45). While urine is a medium that sustains microbial growth, it is well recognised to be nutritionally weak when compared to normal laboratory growth conditions (46). Therefore, it could be argued that uro-associated *E. coli* completely switching off flagellin synthesis to evade TLR5 recognition and the host immune response is not compatible with its survival. However, exploiting population heterogeneity to regulate environmental flagellin concentrations ensures microbial survivability and potentially host colonisation. Additionally, the concept that immune evasion i.e. host TLR5 recognition of flagellin proteins drives uro-associated *E. coli* to downregulate flagellar production may help unravel the pathogenesis of asymptomatic bacteria, defined as the presence of bacteria in the urinary tract without inflammatory symptoms.

It was interesting, however, that following *in vitro* culture the motility and NFκB observations (Fig 2B) did not differentiate between the ASB and cystitis strains. During active UTIs in humans it has been reported that infecting bacteria need to divide rapidly to survive the host innate response. Doubling times have been estimated to be between 17 to 34 minutes and averaging 22 minutes (47). This rapid growth response characterised by upregulation of UPEC translational machinery results in high cell densities that are orchestrated to both overwhelm and escape the host defences (48). The latter is supported by growth experiments using steady state chemostat cultures where faster growing *E. coli* produce more flagella (Sim 2017).

However, a key question relates to the cues in the urogenital tract that trigger increased growth and UPEC infections. It is generally accepted that low nutrient conditions up-regulate *E. coli* flagellar synthesis via activation of the *flhDC* operon (49) although other signals including urine osmolality and pH cannot be ignored (50). Studies in *Salmonella* grown in low nutrient conditions have shown that non flagellar regulators such as RflP, a regulator that modulates ClpXP recognition of FlhD_4_C_2_, can impact the master regulator FlhD_4_C_2_ activity and hence flagellin synthesis (39,44,49,51). Whether comparable regulators function to trigger flagellar growth in uro-associated *E.coli* is not known although NarL, ModE, Metj, GadE and YdeO, all sensors of environmental cues, have been identified as playing potential roles in infection-specific UPEC gene expression (48).

Population heterogeneity is not an original concept and has been shown to be exploited by a number of bacterial species to retain a selective advantage particularly during growth in specific environmental niches (41,52,53). However, its exploitation by uro-associated *E. coli* to regulate flagellin synthesis and avoid the host defences is novel. This study was not designed to identify the regulatory mechanisms functioning to control flagellin synthesis in the urogenital tract, but environmental and genetic cues including population densities, urine osmolality and electrolytes, urinary and bacterial metabolites, and pH need to be investigated further.

## Materials and Methods

### Strains and General Microbiology

*E. coli* strains used in the study have either been previously described (26). Strains PYL3398 to ABU3710 were a kind donation from the Diagnostic Microbiology Unit at the Freeman hospital, Newcastle NHS Trust, Newcastle upon Tyne between 2010 and 2012 (Table S1 and S2). No ethics were necessary for the use of these strains as the researchers did not have access to clinical records and the only information provided by the unit was the type of UTI associated with each isolate. Strains ABU4738-ABU4745 came from the clinical study of Drage et al (2019) that was conducted under ethically approved study protocols (ref: REC-14-NE-0026).

Strains used during this study were propagated in or on Luria-Bertani (LB) medium using 1.5% agar for plates. Incubation, unless stated otherwise, was overnight at 37°C with liquid cultures aerated by orbital shaking at 160 rpm. All motility assays were performed by either direct inoculation using a toothpick or inoculating 3 μl of an overnight culture onto motility agar (1% Tryptone, 0.5% NaCl, 0.3% Agar) and incubating for 8 hours at 30°C. Images of motility swarms were digitally captured, and the vertical and horizontal diameter measured to generate an average swarm distance using ImageJ. All swarm assays were performed with a minimum of three independent colonies. Transformation of the plasmids p*flhDC* or pBAD*flgE*A240C were performed by electroporation as described previously (20). Selection for plasmids was performed using either 100 μg/ml Ampicillin or 50 μg/ml Kanamycin. p*flhDC* was generated by cloning a PCR product using the primers *flhD*-42FBam [ggcggatccGGGTGCGGCTACGTCGCAC] and *flhC*+616RBam [ggcggatccCGCTGCTGGAGTGTTTGTCC] into the high copy number vector pSE280 using standard cloning techniques.

### Isolation of Flagellin and Outer membranes

All flagellin and outer membrane (OM) preparations were based on 1 L cultures of strains grown to an OD_600_ of 0.6-0.7. Cells were centrifuged at 3890 *g* and cell pellets resuspended in cold 10mM HEPES pH 7.4. For flagellin isolation, cell suspensions were sheared using an Ultra-Turrax blending stick for 2 minutes set at 13500 rpm. The same protocol was used prior to OM isolation to reduce flagellin contamination. Blended supernatants were centrifuged at 100,000 g for 1 hour at 4°C to collect sheared flagellar filaments. The pellets were washed by repeating this procedure three times. Pellets were resuspended in 10mM HEPES pH 7.4 and centrifuged at 3890 *g* to improve the removal of cell debris between each ultra-centrifuge wash step. The washed flagellin pellets were resuspended in 500 μl 10mM HEPES pH 7.4 and stored at −20°C.

For outer membrane isolation cell suspensions were lysed using a Constant Systems cell disruptor at 23kPSI. Lysed cell suspensions were centrifuged at 12000 *g* at 4°C for 40 minutes, the supernatant layered onto a sucrose gradient and centrifuged at 56000 *g* for 36 hours at 4°C. Outer membrane fractions were resuspended and washed once in 10mM HEPES pH7.4. The washed outer membrane fraction was collected by centrifuging at 134000 *g* for 6 hours at 4°C and the resulting pellet resuspended in 500 μl 10mmM HEPES. The quality of preparations was assessed using standard SDS polyacrylamide gel electrophoresis.

### NFκB reporter assay, IL-8 ELISA and Immunoblots

Growth conditions for the bladder RT4 cell line has been previously described (26). All challenges were performed using 24 well plates seeded with 500,000 cells in 500 μl, and the cells grown until 80-90% confluent. Bladder RT4 cells were challenged in triplicate with either isolated outer membrane preparations (100 ng/ml protein content), flagellin (0-250 ng/ml) or heat-killed whole cells at 37°C and 5% CO_2_ (26). Challenges were stopped after 24 hours, the extracellular media collected and stored at −20°C. Interleukin 8 concentrations (pg/ml) were assayed using an eBioscience IL-8 ELISA kit following the manufacturer’s instructions. Measurement of RT4 NFκB reporter activity was as previously described (26). For IκBα immunoblots, challenged RT4 cells were lysed in RIPA Buffer collected, quantified using a Micro BCA protein assay kit (Thermo) and either stored at −20°C or 10 μg used for immunoblots with an Anti-IkBa antibody (New England Biolabs) (54).

### Inhibition of TLR4 and TLR5 siRNA expression

TLR4 and TLR5 knockdown experiments using RT4 cells and siRNA were performed as described previously (55). siRNAs used were as follows: s14196 (TLR4) and s14199 (TLR5) and AM4611 (negative siRNA #1) as a control. All challenges were performed as described previously and media bathing the cells analysed using an eBioscience IL-8 ELISA kit following the manufacturer’s instructions.

### Quantification of Flagellar abundance

Expression of *flgEA240C* was analysed following bacterial growth at 37°C in LB media containing 0.1% arabinose with shaking until an OD600 of 0.6 to 0.7. Staining of the cells was performed using AlexaFluor 568 (20). Essentially, bacterial cell suspensions were immobilised on a 1% agarose padded microscope slide and samples analysed in triplicate at 100x objective using a Nikon Eclipse Ti inverted microscope capturing both phase contrast (100 ms exposure) and red channel images (1000 ms exposure) at five different fields of view. Five randomly chosen fields were analysed manually using the ImageJ cell counter plugin generating data where n = 200 – 300 cells. Foci per cell was averaged across the five fields of view to enumerate the level of cell flagellation, as well as the distribution of flagella over the population.

### Data Analysis and Presentation

Data and statistical analysis were performed using MS Excel including the use of ANOVA. Images for figure panels were processed and cropped using ImageJ and imported into Adobe illustrator for formatting. All figures were collated in Adobe illustrator to achieve the correct resolution.

## Acknowledgements

The authors would like to express thanks to Prof John Perry and Dr Kathy Walton of the Diagnostic Microbiology Unit at the Freeman Hospital for the kind gift of *E. coli* isolates used in this study. Finally, we thank Dr James Connolly for his valuable feedback on draft versions of this manuscript.

---

**Table S1:** Motile Strains and associated data used in this study.

| Lab Strain ID | Source* | Flagellin Serotype | Average swarm diameter (cm) | Flagellar abundance § |  | Reference |
| --- | --- | --- | --- | --- | --- | --- |
|  |  |  |  | Avg Fla / cell | % Fla<sup>+</sup> |  |
| Strains used in Figure 1, 2 and 3 |  |  |  |  |  |  |
| 2743 | NCTC10418 |  | 5.78 | 2.35 | 82 | (26) |
| 3373 | CFT073 | H1 | 7.02 | 1.59 | 58 | (56) |
| 3398 | PYL | H1 | 3.59 | 1.46 | 7 | This study |
| 3406 | ABU | H18 | 3.06 |  |  |  |
| 3408 | UTI | H1 | 5.71 | 1.66 | 25 |  |
| 3409 | UTI | H5 | 1.87 |  |  |  |
| 3411 | UTI | H6 | 7.78 |  |  |  |
| 3412 | UTI | H18 | 1.49 | 1.36 | 27 |  |
| 3414 | UTI |  | 7.39 |  |  |  |
| 3415 | BAC | H5 | 3.04 |  |  |  |
| 3417 | UTI | H1 | 2.05 |  |  |  |
| 3419 | ABU | H6 | 3.56 |  |  |  |
| 3424 | PYL | H18 | 4.98 |  |  |  |
| 3425 | PYL | H1 | 4.51 |  |  |  |
| 3692 | ABU |  | 3.07 |  |  |  |
| 3693 | ABU |  | 2.67 | 1.24 | 8 |  |
| 3694 | ABU |  | 5.41 |  |  |  |
| 3695 | ABU |  | 3.96 | 1.41 | 10 |  |
| 3697 | ABU |  | 2.44 | 1.60 | 6 |  |
| 3698 | ABU |  | 2.36 |  |  |  |
| 3699 | ABU | H4 | 0.77 |  |  |  |
| 3701 | UTI | H27 | 2.80 | 1.68 | 52 |  |
| 3702 | UTI | H4 | 6.97 | 1.52 | 24 |  |
| 3703 | UTI |  | 1.00 | 1.71 | 10 |  |
| 3704 | UTI |  | 2.10 |  |  |  |
| 3710 | ABU | H4 | 2.85 |  |  |  |
| Additional strains used in Figure 3 |  |  |  |  |  |  |
| 4738 | ABU | H5 | 1.58 | 1.67 | 50 | (8) |
| 4739 | ABU | H1 | 2.41 | 1.62 | 35 |  |
| 4740 | ABU | H1 | 2.25 | 1.44 | 57 |  |
| 4741 | ABU | H18 | 2.10 | 1.81 | 11 |  |
| 4742 | ABU | H18 | 2.25 | 1.14 | 19 |  |
| 4743 | ABU | H5 | 3.67 | 1.62 | 36 |  |
| 4744 | ABU | H18 | 1.67 | 1.45 | 46 |  |
| 4745 | ABU | H6 | 2.00 | 1.14 | 7 |  |

§ Data shown to indicate the strains used in Figure 3
\*PYL: Pyelonephritis; ABU: asymptomatic bacteriuria; UTI: Acute cystitis; BAC:
Bacteraemia

**Table S2.** Non-motile strains used in Figures 1 and 2D.

| Lab Strain ID | Source* |
| --- | --- |
| 3399 | ABU |
| 3400 | ABU |
| 3401 | UTI |
| 3402 | BAC |
| 3403 | UTI |
| 3407 | UTI |
| 3410 | BAC |
| 3413 | ABU |
| 3416 | ABU |
| 3418 | ABU |
| 3420 | ABU |
| 3422 | PYL |
| 3696 | ABU |
| 3701 | UTI |
| 3706 | UTI |
| 3707 | UTI |

\*See Table S1 descriptors

